# A Saliva-Inspired Model of Enzymatic Starch Hydrolysis by Alpha Amylase

**DOI:** 10.64898/2026.11.03.764040

**Authors:** Rajni Kudawla, Sourav Negi, Avneesh Singh, Deepika Singh, Tripta Bhatia

## Abstract

We demonstrate that the enzymatic activity of alpha-amylase (AA) can serve as an effective physicochemical trigger for membrane remodeling in giant unilamellar vesicles (GUVs). This work provides insight into how primitive protocellular reactions directly translate into the membrane’s mechanical responses. Analysis of the potato starch utilized in this work is composed of 26% amylose and 75% amylopectin. Quantitative DNS analysis revealed time-and concentration-dependent accumulation of reducing sugars during starch degradation by AA. The HPLC-RID chromatograms demonstrated the synthesis of glucose, maltose, maltotriose, maltotetraose, maltopentaose, maltohexaose, and maltoheptaose, with maltose and maltotriose as the major products. The accumulation of these soluble hydrolysis products gradually increases the osmolarity of the extracellular medium surrounding the vesicles, leading to osmotic asymmetry across the membrane. This biochemical framework provides a quantitative basis for understanding membrane deformation, curvature generation, vesicle deflation, and budding. In addition to the specific starch–***α***-amylase system studied here, our work uncovers a general mechanism by which external biochemical reactions control membrane shape transformations via osmotic coupling. Such enzyme-driven remodeling offers a simple yet powerful strategy for inducing dynamic structural changes in artificial cells. It may provide insight into primitive protocellular processes in which metabolic activity and membrane mechanics were inherently coupled.

## 1 Introduction

In the last decades, human saliva has been widely studied, especially because of its promise as a non-invasive diagnostic and prognostic medium. The identification of its diverse molecular components and physicochemical properties has led to the development of saliva-based techniques for monitoring different diseases [1–4]. Human saliva is a complex biological fluid that is made up of a range of organic and inorganic substances. In addition to water and electrolytes, saliva contains proteins and glyco-proteins, enzymes, immunoglobulins, hormones and lipids [5, 6]. Neutral and polar lipids have previously been found in human saliva, such as cholesterol, triglycerides, free fatty acids, phospholipids and glycolipids [7, 8]. However, the reported quantities are highly variable depending on the salivary gland studied and the method of sample collection and lipid analysis. Total lipids in entire saliva have been reported to be approximately 1.3 mg/100 ml with neutral lipids being more than 95% of the total lipid fraction [5]. Conversely, it has been found that secretions from small salivary glands contain relatively significant levels of polar lipids, such as phospholipids and glycolipids. Phospholipids represent around 3.6% of the total lipid fraction of entire saliva, but their proportional amount varies in different salivary discharges [5, 9]. The results suggest that lipids are present in the salivary milieu, but their biological activities and interactions with other salivary elements are not fully understood. Saliva also contains the digestive enzyme *α*-amylase to commence the enzymatic breakdown of ingested starch in the mouth.

Starch is a primary carbohydrate in the diet and a key source of metabolic energy [10]. It is primarily composed of two macromolecular fractions, amylose and amylopectin [11]. Amylose is largely a linear polymer of D-glucose units linked through *α*-(1*→*4) glycosidic bonds, whereas amylopectin is a highly branched polysaccharide containing *α*-(1*→*4)-linked chains interconnected through *α*-(1*→*6) glycosidic linkages. Because of this structural complexity, starch degradation depends on the coordinated activity of multiple enzymes collectively referred to as amylases [12–15]. Among starch-degrading enzymes, *α*-amylase (EC 3.2.1.1) is of particular importance as an endo-acting hydrolase that cleaves internal *α* (1→4) glycosidic bonds within starch chains. Its action generates smaller oligosaccharides such as maltose, maltotriose, and *α*-limit dextrins [11, 16–18], thereby gradually changing the makeup of the aqueous environment around. The rate and extent of starch digestion are important determinants of postprandial metabolic responses, making the biochemical processing of starch relevant to metabolic health [19–21].

The biochemical aspects of starch hydrolysis by *α*-amylase have been extensively studied, but substantially less attention has been paid to the possible physical effects of this enzymatic activity on lipid membrane systems. This question is especially interesting since biological membranes are active structures whose shape and mechanical properties may adapt to changes in their chemical and osmotic surroundings. Additionally, the presence of lipids in saliva raises the potential that interactions between products of starch digestion and lipid-containing compartments can impact membrane behaviour.

To answer this question in a controlled manner, we have constructed a simple saliva-inspired model system based on potato starch, malt-derived *α*-amylase and lipid membranes. Malt amylase is a plant-derived amylase that catalyses the hydrolysis of starch and therefore provides a convenient experimental setup for investigating the physical consequences of starch degradation. Importantly, previous physiological studies have directly compared malt amylase with human salivary amylase in starch digestion. The early rates of starch digestion produced by the two enzymes were reported to be similar, while malt amylase was able to remain active for a longer period under acidic gastric conditions. The study also showed that malt amylase could substantially enhance starch digestion under conditions in which salivary amylase was restricted by acid inactivation. Therefore, these observations demonstrate that malt amylase is a functionally relevant model enzyme for the investigation of starch hydrolysis, avoiding the complexity and variability associated with the use of human saliva [22]. Phospholipid Giant unilamellar vesicles (GUVs) were used as membrane compartments, with cholesterol incorporated into selected membrane compositions to examine the influence of membrane composition on the response to enzymatic starch hydrolysis. GUVs are cell-sized vesicles enclosed by a single lipid bilayer and provide a well-controlled model system for investigating membrane biophysics [23, 24]. Their micron-scale size also allow direct visualisation of membrane morphology and deformation using optical microscopy [25].

In the present study, we characterise *α*-amylase-mediated starch hydrolysis using biochemical 3,5-dinitrosalicylic acid (DNS) assays and high-performance liquid chromatography (HPLC) and investigate the resulting response of phospholipid and cholesterol-containing GUVs. By combining biochemical characterisation of starch hydrolysis with quantitative visualisation of membrane morphology, we aim to establish a connection between enzymatic starch digestion and the physical remodelling of lipid membrane compartments. This simplified model provides a framework for investigating how biochemical reactions occurring in a saliva-like environment can be coupled to membrane-level physical responses.

## 2 Materials and Methods

### 2.1 Materials

1-palmitoyl-2-oleoyl-sn-glycero-3-phosphocholine (POPC) lipid (Catalog number 850457P) and Cholesterol (Catalog number C8503) were purchased from Avanti Polar Lipids in the form of powder and dissolved in chloroform to make a stock solution of 1 mg/ml. Lipid stock solutions are stored at *−*20*^◦^C*. Starch soluble GR (CAS No. 9005-84-9), Amylose from potato (CAS No. 9005-82-7), and amylopectin from maize (CAS No. 9037-22-3) and for confocal experiments nile red dye (Catalog number 7385-67-3) and Fluorescein isothiocyanate (FITC) dye (Catalog number 27072-45-3) were obtained from Sigma-Aldrich. Alpha-amylase from Malt was purchased from HI media (Catalog number 70 GRM638). For the enzymatic hydrolysis assay, KOH (from SRL, CAS no. 1310-58-3), HCl about 37% for analysis (CAS No. 7647-01-0), Potassium Sodium Tartarate (CAS no. 6381-59-5) also known as Rochelle salt, 3,5-Dinitrosalicylic acid (CAS no. 609-99-4) and Iodine (CAS no. 7553-56-2), all were purchased from sigma merch aldrich. For the HPLC experiment, glucose (CAS No. 50-99-7) from Sigma-Aldrich, dextrin and all maltooligosaccharides standards were purchased from BLDpharmatech India Pvt Ltd. HPLC-grade acetonitrile was procured from J. T. Baker. Milli-Q ultrapure water, having resistivity of 18.2 MΩcm at 25*^◦^C*, was used throughout the experiment. All the solutions for HPLC experiments were prepared in ultrapure water unless otherwise stated. It served both as the reaction medium and as the diluent for all stocks and standards.

### 2.2 Methods

#### 2.2.1 Determination of Amylose/Amylopectin Content

The amylose and amylopectin contents of potato starch were quantified using an iodine-binding spectrophotometric assay [26, 27]. Briefly, 5 *mg* of starch powder was weighed into a small beaker and dissolved in 2 *mL* of 0.5 *N* KOH. The mixture was heated at 60*^◦^C* to 70*^◦^C* under magnetic stirring for 5–10 min to ensure complete dispersion. The resulting solution was transferred to a 10 *ml* centrifuge tube and diluted to a final volume of 10 *ml* with deionized water.

An aliquot (0.2 *ml*) of this stock solution was then transferred to a fresh 10 *ml* tube, neutralized with 0.5 *ml* of 0.1 *N* HCl, and reacted with 0.1 *ml* of iodine reagent. The final volume was adjusted to 5 *ml* with deionized water, and the absorbance of the developed blue complex was measured at 625 *nm* using a UV–Vis spectrophotometer. Amylose content was determined from a calibration curve prepared using known mixtures of amylose and amylopectin standards. For each standard, accurately weighed amounts of amylose and amylopectin were combined such that the total mass was 5 *mg*. The mixture was dissolved in 2 *ml* of 0.5 *N* KOH, thoroughly mixed until completely solubilized, and diluted to 10 *ml* with distilled water. These standards were processed in the same manner as the samples for absorbance measurement.

#### 2.2.2 Enzymatic hydrolysis of starch

The extent of starch hydrolysis in the external medium was quantified using the 3,5-dinitrosalicylic acid (DNS) colorimetric assay [27, 28] to determine reducing sugar concentration as a function of time. All enzymatic reactions were performed under conditions identical to those used during vesicle imaging experiments to ensure direct correlation between biochemical activity and membrane morphology. Starch solution (0.5 mg/mL) was prepared in the same buffer system used for GUV experiments. Hydrolysis was initiated by adding defined concentrations of *α*-amylase (1 *−* 3 *µM*) to the starch solution at room temperature. The reaction mixtures were gently mixed and incubated under static conditions. At predetermined time intervals, aliquots were withdrawn for reducing sugar analysis. To terminate the enzymatic reaction, 200 *µl* of the reaction mixture was immediately mixed with 600 *µl* of DNS reagent (sample: DNS ratio = 1:3). The reaction mixture was heated in a boiling water bath for 10 min to allow color development and then cooled to room temperature. The samples were centrifuged at 2000 rpm for 5 min to remove suspended particles. However, due to the low starch concentration (0.5 mg/mL), no visible pellet was observed after centrifugation; therefore, the entire supernatant was used for absorbance measurement at 540 *nm* using a UV–Vis spectrophotometer. For spectrophotometric measurements, 200 *µ*L of the hydrolysed starch reaction mixture was diluted to a final volume of 3 mL in the cuvette before absorbance measurement at 540 nm. A calibration curve was constructed using glucose standards (see Supporting Information, Figure S3), and reducing sugar concentrations were determined from the linear regression equation. The degree of hydrolysis was expressed as dextrose equivalent (DE%), calculated as the percentage ratio of reducing sugars (expressed as glucose equivalents) to the initial starch concentration. The dextrose equivalent (DE%) was calculated using the following relation:

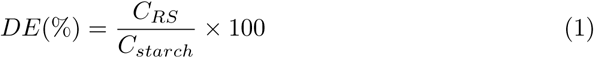

where *C_RS_*represents the concentration of reducing sugars (expressed as glucose equivalents) determined from the glucose calibration curve, and *C_starch_* denotes the initial starch concentration present in the reaction mixture.

#### 2.2.3 HPLC-RID Analysis of Starch Hydrolysis Products

The hydrolysis products obtained by *α*-amylase degradation of potato starch were analysed by HPLC with a refractive index detector (RID). For this, a starch stock solution of 0.625 mg mL*^−^*^1^ and an *α*-amylase stock solution of 17.68 *µ*M were prepared separately. The reaction mixtures used for enzymatic hydrolysis and subsequent HPLC analysis were prepared according to Table 1.

**Table 1.** Composition of reaction mixtures used for enzymatic hydrolysis of potato starch at different *α*-amylase concentrations for HPLC analysis.

| **α-amylase<br>concentration (µM)** | **Starch solution<br>(µL)** | **α-amylase<br>(µL)** | **Ultrapure water<br>(µL)** | **Total volume<br>(µL)** |
|---|---|---|---|---|
| 1.0 | 8000 | 560 | 1440 | 10000 |
| 1.5 | 8000 | 840 | 1160 | 10000 |
| 2.0 | 8000 | 1120 | 880 | 10000 |
| 2.5 | 8000 | 1400 | 600 | 10000 |

After 30 min incubation, 1 mL aliquot was removed from each reaction mixture and the enzymatic reaction was stopped by heating the sample at 100 *^◦^*C for 2 min. Samples were then kept on ice and centrifuged at 7500*×g* for 5 min to remove insoluble residue. Because of the low concentration of hydrolysis products, the quenched samples were concentrated with a vacuum concentrator at 45 *^◦^*C until all solvent was removed. The residue was dissolved in ultrapure water and utilised for HPLC analysis.

Chromatographic separation was carried out on an aminopropyl (NH_2_) column (250 *×* 4.6 mm, 5 *µ*m) with a guard column of the same kind. The mobile phase was a combination of acetonitrile and water (70:30, v/v) under isocratic conditions with a flow rate of 0.8 mL min*^−^*^1^. The column temperature was maintained at 30 *^◦^*C and the refractive index detector at 35 *^◦^*C. The injection volume was 10 *µ*L and the entire time of the run was 40 min.

Hydrolysis products were identified by comparing retention periods to commercially available standards of glucose, maltose, maltotriose, maltotetraose, maltopentaose, maltohexaose and maltoheptaose (see Figure 5). The chromatographic profiles of the hydrolysed starch samples were utilized to elucidate the nature of the oligosaccharides generated during the breakdown of starch by *α*-amylase.

#### 2.2.4 Preparation of Giant Unilamellar Vesicles (GUVs)

Two different types of GUVs composed of only 1-palmitoyl-2-oleoyl-sn-glycero-3-phosphocholine (POPC) and POPC: Cholesterol (9:1) were prepared using the electroformation method described by Angelova and Dimitrov [29]. These GUVs were prepared in 0.5 mg/mL starch solution at a temperature of 37*^◦^*C as shown in Figure 1. For the experiments, stock solutions of starch and *α*-amylase, each at a concentration of 0.5 mg/ml, were prepared in Milli-Q water. The electroformation procedure followed in this study was adapted from our previously reported protocol [30].

**Fig. 1.**
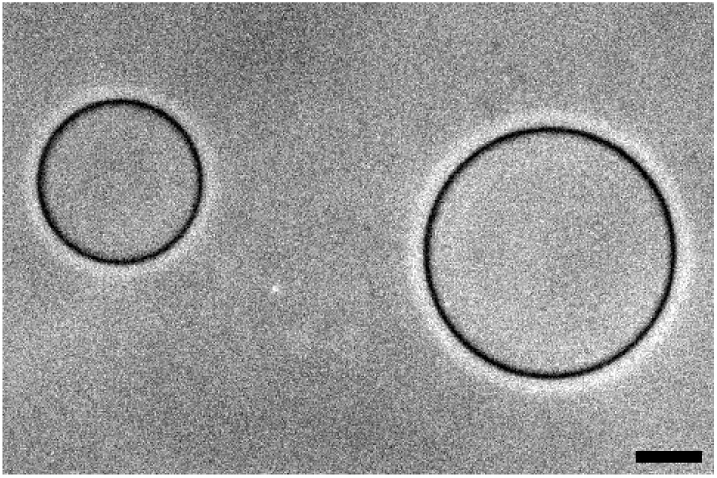
Phase contrast image of two giant unilamellar vesicles prepared in 0.5 mg/ml starch. The black contours represent the lipid bilayer. There is a starch solution inside and outside the membrane bilayer. The scale bar for the image is 10 *µm*.

#### 2.2.5 Preparation of Buffer Solution

For preparing a stock solution of starch in milli-Q water, weigh 5 *mg* of soluble starch powder and add 7 *−* 8 *ml* of warm milli-Q water to the starch powder. Heat the starch solution from 70*^◦^C* to 80*^◦^C* while stirring with a magnetic stirrer at 500 rpm until the starch is fully dissolved. Finally, top up to 10 *ml* in falcon tube with milli-Q water. Leave this falcon tube in Rota spin for at least 5-10 minutes for mixing well. Now, this starch solution [31] is ready to make GUVs. The fresh buffer solution was prepared for the GUV preparation throughout the experiments. Different concentrations of *α*-amylase used in the experiments were prepared using a calculated amount of *α*-amylase from the 0.5 mg/mL (8.93 *µM*) stock solution prepared before.

#### 2.2.6 Phase-Contrast and Confocal Microscopy of GUV Remodeling

##### Microscopy conditions

Vesicle imaging was performed using phase contrast microscopy on a Zeiss Axio Observer 5, which was equipped with a 20x objective (PL APO 20x/0.8 Ph2 M27, working distance 0.5 *mm*). A Zeiss 506 mono camera was used to acquire images and time-lapse videos. Time-lapse phase-contrast videos were acquired to monitor the temporal evolution of GUV morphology following the addition of *α*-amylase. Illumination was provided by a transmitted light (TL) LED source. The exposure time was fixed at 29 ms throughout the experiments. All videos and images were analysed using ImageJ software.

##### Fluctuation data collection

To determine the mechanical properties of POPC GUVs in a 0.5 mg/mL starch solution in the absence of *α*-amylase, vesicle fluctuation (flicker) spectroscopy was performed following established methods [32]. Thermally fluctuating vesicles were recorded using the same microscope setup described in the microscopy section above. For each vesicle, approximately 10000 frames were acquired to ensure sufficient statistical sampling of membrane fluctuations. Image sequences were recorded at an exposure time corresponding to 29 frames per second, allowing accurate capture of the thermally driven contour fluctuations used for the subsequent bending rigidity analysis.

##### Nile Red imaging

Nile Red was used as a membrane fluorescent probe to visualize the lipid bilayer and the morphological transformations of GUVs. Confocal fluorescence images and *z*-stacks were acquired to characterize membrane morphology and the formation of internal buds and compartments.

##### Fluorescent labeling of *α*-amylase

To look at the interaction of *α*-amylase with the membrane and investigate its pathway of entry into the membrane, *α*-amylase was labelled with FITC dye [33]. For labelling, prepare a stock solution of 1 mg mL*^−^*^1^ FITC dye in DMSO (prevent FITC from direct light), 0.1 M sodium carbonate buffer at pH 9, and 1 M ammonium chloride buffer. Mix 450 *µ*L of sodium carbonate buffer (0.1 M) with 3 mg *α*-amylase. Add 50 *µ*L of 1 mg mL*^−^*^1^ FITC dye into that solution. Leave this mixture in the dark for 5–6 h at 4 *^◦^*C by covering using silver foil. After 5–6 h, add 50 *µ*L of ammonium chloride buffer to this mixture to quench the dye and leave this MCT for 3–4 h at 4 *^◦^*C. The unbound FITC from the conjugate was separated using the sephadex column in different MCTs. Then, using a UV-Vis spectrometer, check the absorbance of protein and dye as shown in Figure 2. The simultaneous presence of both peaks confirms the successful conjugation of FITC with the enzyme.

**Fig. 2.**
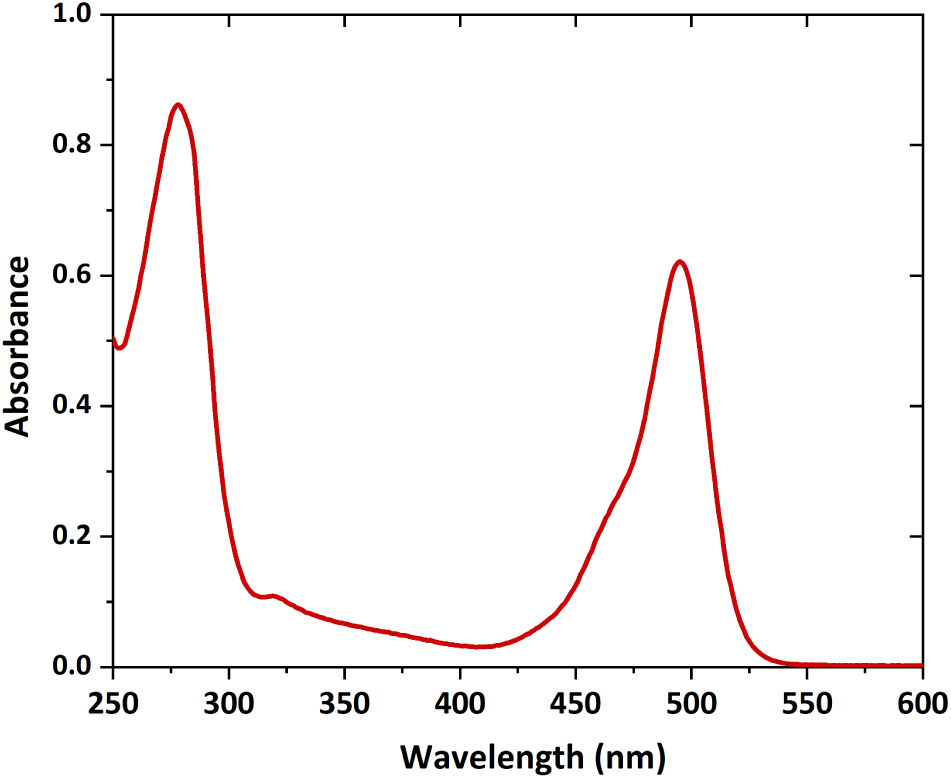
UV–Visible absorption spectra of FITC-labeled *α*-amylase after purification through a sephadex column. The characteristic absorption peaks observed near 280 nm and 495 nm correspond to *α*-amylase and FITC dye, respectively.

## 3 Results

### 3.1 Compositional Characterization

The potato starch used in this study consisted of 25.7% amylose and 74.3% amylopectin [18, 34], as determined by the iodine-binding spectrophotometric method (see Supporting Information, Figures S1 and S2). This composition is consistent with typical potato starch, which is dominated by highly branched amylopectin chains with a smaller fraction of linear amylose. The relative abundance of amylopectin is particularly relevant for enzymatic degradation, as its branched architecture provides multiple cleavage sites for *α*-amylase while simultaneously influencing the evolution of molecular weight distribution during hydrolysis. The presence of 25.7% amylose introduces linear chains that may contribute differently to osmotic behaviour and degradation kinetics. Some features of amylose and amylopectin are mentioned in Table 2.

**Table 2.** Comparison of amylose and amylopectin, the two major polysaccharide components of potato starch used in this study.

| **Features** | **Amylose** | **Amylopectin** |
|---|---|---|
| Structure | Linear polymer | Highly branched polymer |
| Glycosidic linkage | α(1 → 4) | α(1 → 4) & α(1 → 6) |
| Branching | unbranched mostly | Branch points every 24–30 glucose units |
| Molecular size | Smaller molecular weight | Very large molecular weight |
| Content in starch | 26% | 75% |
| Solubility | Less soluble | More soluble |

### 3.2 Quantification of Starch Hydrolysis by DNS Assay

To probe the biochemical activity occurring outside the vesicles, 3,5-dinitrosalicylic acid (DNS) analysis was performed to quantify reducing sugar production in the external medium. The reducing sugar concentration was calculated by dividing the obtained glucose mass by the sample volume. The temporal evolution of reducing sugar concentration at different *α*-amylase concentrations is shown in Figure 3.

**Fig. 3.**
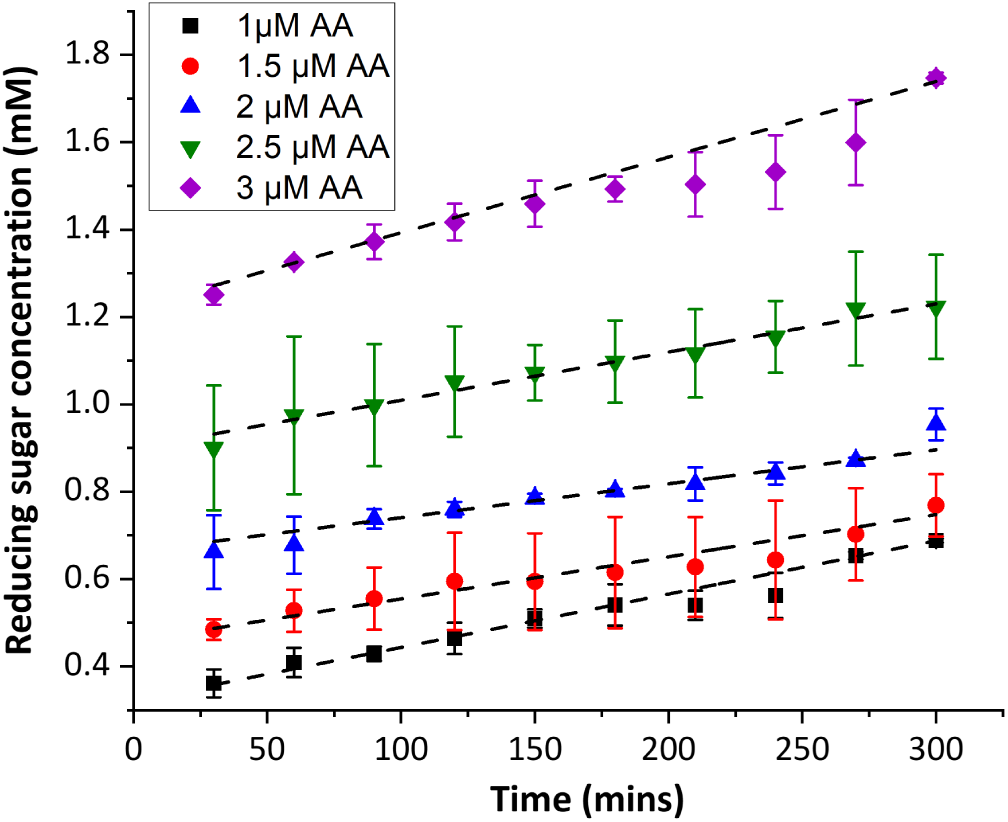
Reducing sugar concentration as a function of time during *α*-amylase-mediated hydrolysis of potato starch at different enzyme concentrations. Symbols denote experimental data, and black dotted lines represent linear regression fits over the interval (30–300 min). The slope of each fit corresponds to the rate of reducing sugar production.

The concentration of reducing sugar increased approximately in a linear fashion with time for all enzyme concentrations studied. This indicates ongoing enzymatic cleavage of *α*-(1*→*4) glycosidic links in starch. The positive slopes recorded for all the data sets confirm the increasing formation of soluble hydrolysis products throughout the reaction period.

The fitted slopes obtained over 30–300 min varied across the *α*-amylase concentrations studied, ranging from 0.000776 *±* 4.55 *×* 10*^−^*^5^ to 0.00173 *±* 5.01 *×* 10*^−^*^5^ mM, min*^−^*^1^. The slope initially decreased with increasing *α*-amylase concentration, from 0.00122 *±* 3.28 *×* 10*^−^*^5^ mM, min*^−^*^1^ at 1 *µ*M to 0.000776 *±* 4.55 *×* 10*^−^*^5^ mM, min*^−^*^1^ at 2 *µ*M, followed by an increase at higher concentrations. The 2 *µ*M *α*-amylase system showed the lowest hydrolysis rate, whereas the 3 *µ*M system exhibited the highest rate (0.00173*±*5.01*×*10*^−^*^5^ mM, min*^−^*^1^). Thus, although the rates did not increase monotonically with enzyme concentration, the higher rate observed at 3 *µ*M suggests enhanced starch hydrolysis at the highest enzyme concentration tested. Overall, the positive slopes for all concentrations indicate continuous enzymatic degradation of starch and progressive formation of reducing sugars in the external medium, generating osmotically active hydrolysis products. The higher rate and extent of hydrolysis at elevated enzyme concentrations indicate enhanced cleavage of *α*-1,4 glycosidic bonds in starch, leading to the accumulation of osmotically active species such as glucose and maltose in the external solution.

The calculated DE% values followed a similar concentration-dependent trend, confirming progressive starch degradation as shown in Figure 4. The DE% increases progressively with reaction time, indicating the gradual conversion of starch into smaller sugar molecules. Higher enzyme concentrations result in a faster increase in DE%, confirming enhanced catalytic activity of AA.

**Fig. 4.**
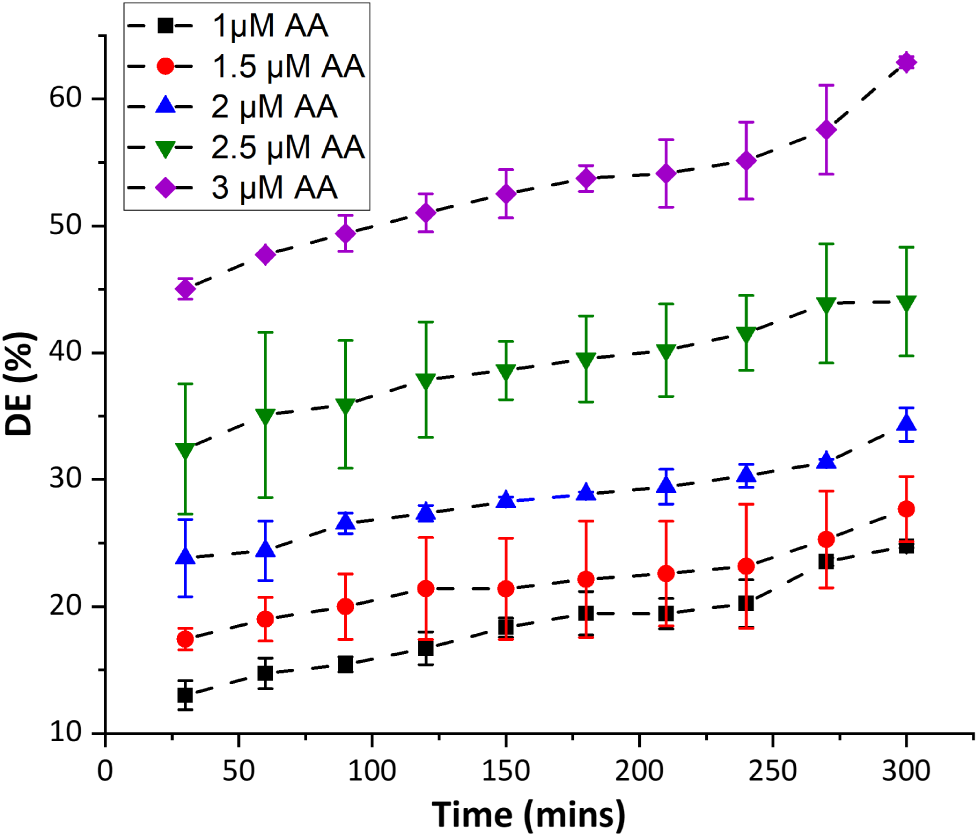
Time-dependent change in dextrose equivalent (DE%) during *α*-amylase (AA)–mediated hydrolysis of potato starch at varying enzyme concentrations. The increase in DE% indicates the extent of starch degradation and the formation of smaller reducing sugar molecules as enzymatic hydrolysis progresses.

**Fig. 5.**
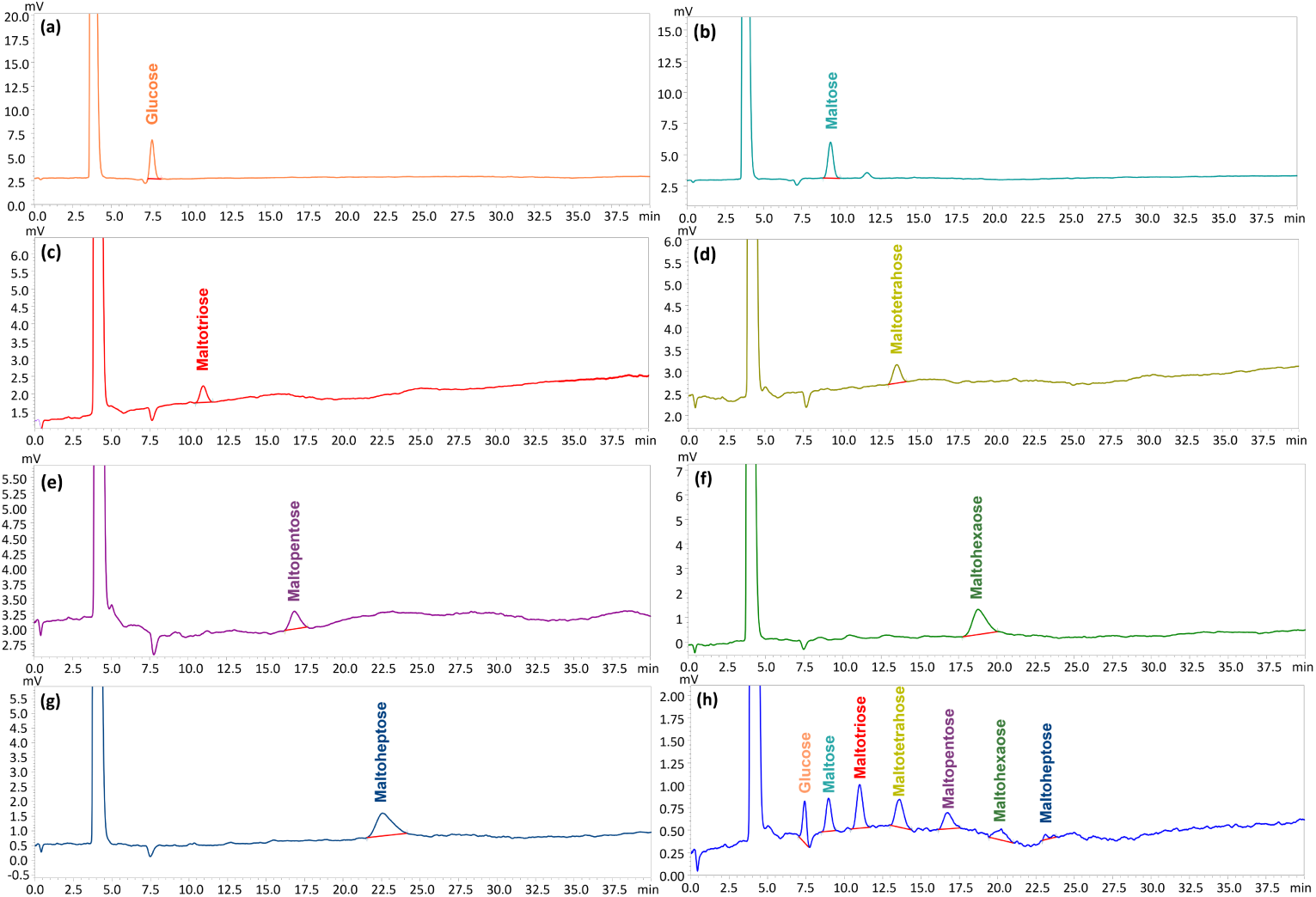
HPLC–RID chromatograms establishing the specificity of the method for individual sugars: (a) glucose (retention time 7.1 min), (b) maltose (8.9 min), (c) maltotriose (10.9 min), (d) maltotetraose (13.5 min), (e) maltopentaose (16.7 min), (f) maltohexaose (19.1 min), and (g) maltoheptaose (23.2 min), and (h) Chromatogram of a standard mixture containing all seven sugars, glucose, maltose, maltotriose, maltotetraose, maltopentaose, maltohexaose, and maltoheptaose at a concentration of 100 *µ*g mL*^−^*^1^.

At the lowest enzyme concentration, 1 *µ*M the DE% increased gradually from approximately 13% at 30 min to about 25% at 300 min. Increasing the enzyme concentration to 1.5 and 2 *µ*M resulted in higher DE values throughout the hydrolysis period, reaching approximately 28% and 34%, respectively, after 300 min. At 2.5 *µ*M, a much larger rise was seen in which the DE% jumped from approximately 32% to nearly 44%. The highest enzyme concentration studied, 3 *µ*M, exhibited the greatest extent of hydrolysis, with the DE% increasing from approximately 45% at 30 min to about 63% at 300 min. These results demonstrate that increasing enzyme concentration enhances the rate and extent of starch depolymerisation.

### 3.3 HPLC Characterization of Hydrolysis Products

Hydrolysed samples were examined by HPLC-RID to determine the compounds formed during enzymatic degradation of potato starch. Individual standards of glucose, maltose, maltotriose, maltotetraose, maltopentaose, maltohexaose and maltoheptaose were chromatographed under identical conditions prior to sample analysis to determine retention times, as shown in Figure 5.

The aminopropyl column produced sufficient separation of the maltooligosaccha-rides based on their molecular size and their interaction with the stationary phase. Under the given chromatographic conditions, each standard produced a distinct and reproducible peak, allowing for reliable assignment of retention durations. The carbohydrate species generated during enzymatic hydrolysis were identified by comparing the retention periods of the hydrolyzed starch samples with those of the standards. Figure 6 shows the chromatograms obtained for the starch hydrolysis reactions performed at different *α*-amylase concentrations after 30 min incubation. In all cases, many peaks attributable to glucose and malto-oligosaccharides were detected. The identification of the products was carried out by comparing the retention durations of the peaks of the samples with those of the reference standards (see Figure 5). The chromatograms indicated that the major hydrolysis products were maltose and maltotriose with minor quantities of glucose and higher malto-oligosaccharides, including maltotetraose, maltopentaose, maltohexaose and maltoheptaose. The existence of these products is compatible with the endo-hydrolytic action of *α*-amylase which hydrolyses internal *α*-(1→4) glycosidic bonds in amylose and amylopectin chains, resulting in a range of soluble oligosaccharides rather than the complete conversion to glucose.

**Fig. 6.**
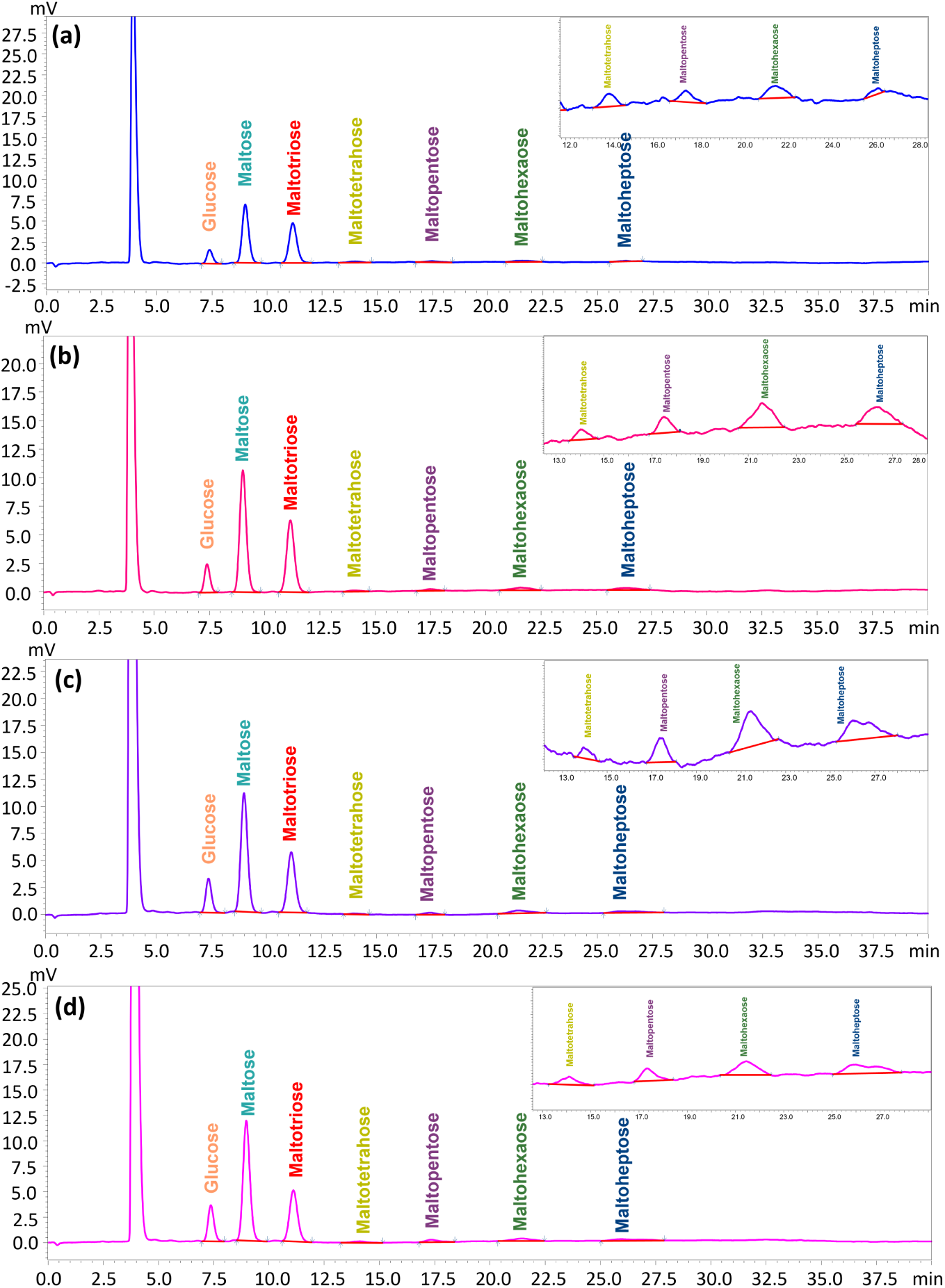
HPLC–RID chromatograms of starch hydrolysates obtained after 30 min of incubation with *α*-amylase at different enzyme concentrations: (a) 1 *µ*M, (b) 1.5 *µ*M, (c) 2 *µ*M, and (d) 2.5 *µ*M. Peaks were identified by comparison with the retention times of glucose and malto-oligosaccharide standards. The inset presents a magnified view of the chromatogram highlighting the low-abundance maltooligosaccharide peaks that are not easily seen in the full-scale chromatogram.

In addition to maltose and maltotriose, maltotetraose, maltopentaose, maltohexaose and maltoheptaose were found in detectable concentrations. The existence of both short and longer oligosaccharides at the same time suggests that the hydrolysis of starch does not happen as a single-step conversion, but as a series of intermediate products. They are produced by the gradual fragmentation of amylose and amylopectin chains into smaller soluble species.

Similar product distribution was observed for all concentrations of enzymes examined, showing that the nature of the hydrolysis products did not alter with concentration in the analysed range. However, the chromatographic peak strength increased with the rise in *α*-amylase concentration, indicating a higher extent of starch degradation and increased generation of soluble sugars at higher enzyme levels.

The insets show the chromatograms magnified, with the low intensity peaks of the higher maltooligosaccharides. The peaks are much smaller than those of maltose and maltotriose, but their systematic occurrence in all samples demonstrates the formation of a broad spectrum of hydrolysis products. The presence of higher oligosaccharides also indicates that after 30, min of incubation, the hydrolysis was not complete, and that a fraction of partially digested starch fragments was still in solution.

The HPLC results thus support the hydrolysis of potato starch by *α*-amylase to a combination of glucose and malto-oligosaccharides, with maltose and maltotriose being the major products. The HPLC-based estimate accounts for the molecular distribution of the soluble hydrolysis products and provides a more composition-specific estimate of their osmotic contribution than the DNS assay. These values represent the estimated osmotic contribution of the externally generated hydrolysis products and should not be interpreted as a direct measurement of the transmembrane osmotic-pressure difference. The latter depends on the osmolarity of both the internal and external solutions and on the permeability of the lipid bilayer to the hydrolysis products.

### 3.4 Alpha-Amylase Induced Remodelling of POPC GUVs

GUVs were prepared in 0.5 mg/mL starch solution to establish an initially symmetric osmotic environment across the membrane as shown in Figure 1. In the absence of *α*-amylase, the vesicles are called “mother vesicles” and contain starch polymer inside and outside. In the absence of external *α*-amylase, vesicles retained a stable spherical morphology over extended observation times, and no detectable shape transformations were observed, confirming osmotic equilibrium and membrane stability under control conditions.

Upon addition of *α*-amylase at varying concentrations to the external medium (defined as t = 0 min), time-dependent morphological changes became evident under phase-contrast microscopy because *α*-amylase starts to break down the long chain of polymer into smaller glucose units, i.e, oligosaccharides of different chain length and alpha-limit dextrins [13]. Oligosaccharides are made up of two or more saccharide units, whereas alpha-limited dextrin contains a branched part. Individual vesicles were monitored continuously to track their structural evolution. Shortly after enzyme addition, localized inward membrane deformations appeared, which progressively developed into distinct internal buds or daughter vesicles as shown in Figure 7.

**Fig. 7.**
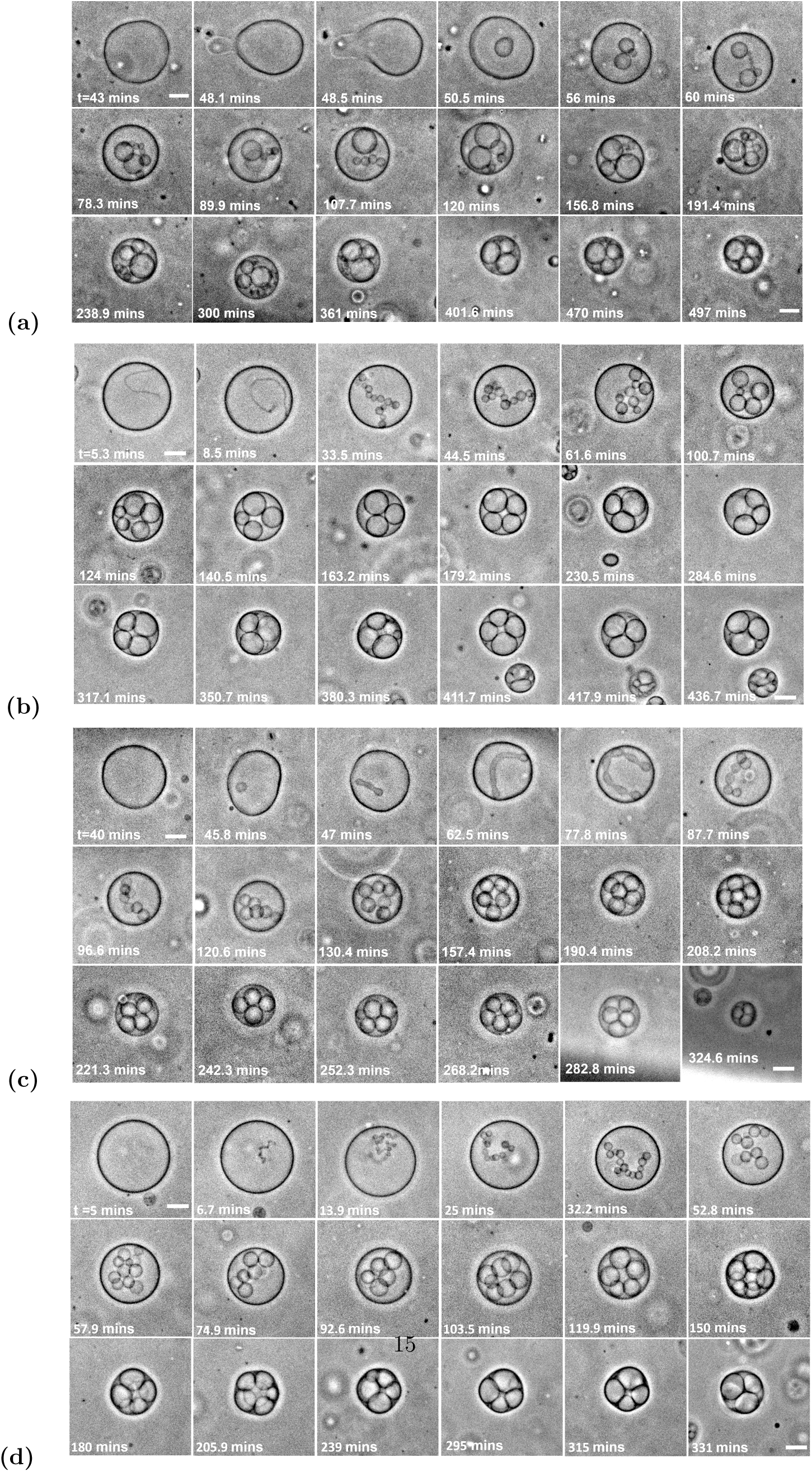
Time-lapse microscopy images showing morphology changes of POPC GUVs (initial diameter: 31–33 *µ*m) in the presence of increasing concentrations of *α*-amylase: (a) 1 *µ*M, (b) 1.5 *µ*M, (c) 2 *µ*M and (d) 2.5 *µ*M. Enzymatic hydrolysis of starch produces soluble sugars that increase the osmotic concentration of the surrounding medium, generating an osmotic pressure difference across the membrane and resulting in progressive membrane deformation. Time zero (*t* = 0 min) corresponds to enzyme addition. Scale bar: 10 *µ*m for all images.

As the concentration of *α*-amylase increases, the rate of inner bud generation also increases, resulting in a reduced interaction time between *α*-amylase and the GUV. At higher enzyme concentrations, vesicle compaction and multi-compartment formation occur within shorter timescales, indicating accelerated osmotic deflation driven by enhanced enzymatic hydrolysis. Over time, these internal buds expanded in size, while the overall diameter of the mother vesicle gradually decreased. Notably, the newly formed internal compartments remained enclosed within the original membrane boundary, indicating that the total bilayer area was conserved, as shown in Figure 8. The reduction in mother vesicle size therefore reflects redistribution of membrane area into inward buds under conditions of decreasing vesicle volume. The projected areas of the mother vesicle and the internal buds were measured from phase-contrast microscopy images using ImageJ by manually tracing the vesicle contours at different time points.

**Fig. 8.**
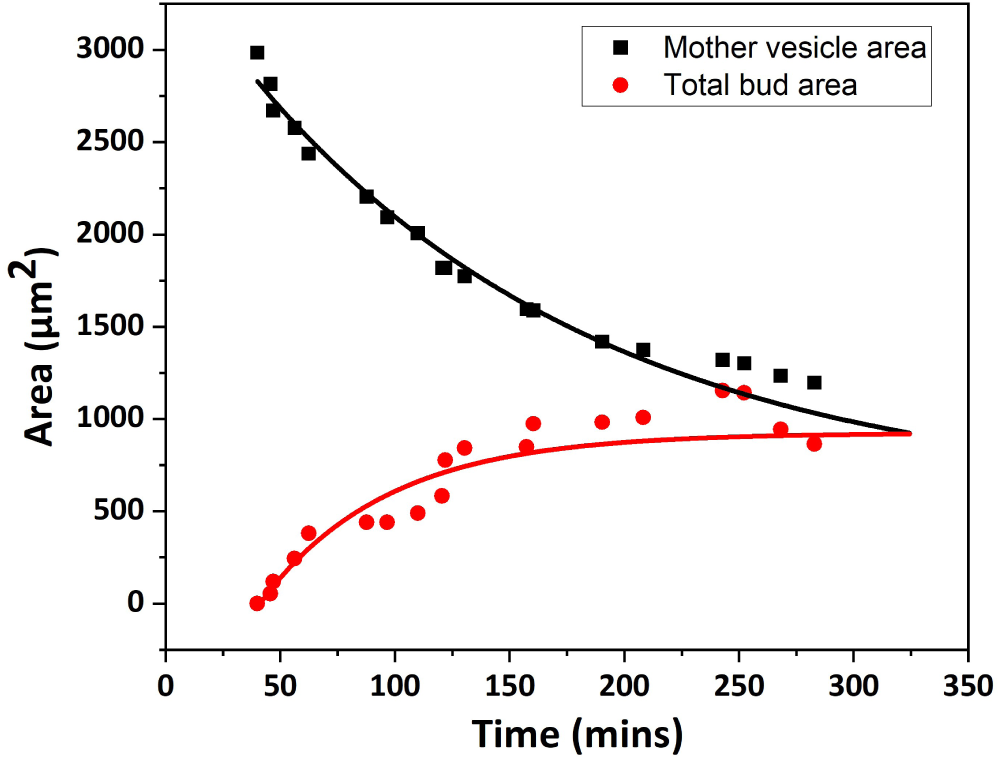
Evolution of mother vesicle and internal bud areas in POPC GUVs exposed to 2 *µ*M *α*-amylase (see Figure 7 c). The mother vesicle area (black, left axis) decreases as the total internal bud area (red, right axis) increases with time. The internal compartments remain enclosed within the original membrane boundary, indicating conservation of the total lipid bilayer area. Solid curves show exponential fits to the data.

As the enzymatic reaction proceeded, budding events became more pronounced and multiple internal compartments were observed. The progressive compaction of the vesicle suggests a time-dependent reduction in reduced volume driven by osmotic imbalance generated in the external medium during starch hydrolysis.

### 3.5 Effect of Cholesterol on GUV Remodeling

Since cholesterol is one of the lipid components reported in human saliva, we next incorporated cholesterol into the model membrane system to better represent the lipid composition considered in our saliva-inspired model. To investigate whether the presence of cholesterol influences the membrane response to enzymatic starch hydrolysis, GUVs composed of POPC: Cholesterol (9:1) were prepared and exposed to different concentrations of *α*-amylase. The morphological response of these cholesterol-containing GUVs was then compared with that of the POPC GUVs described above. Phase-contrast microscopy was used to track the morphological evolution of these vesicles after the addition of alpha-amylase. After *α*-amylase addition, POPC: Cholesterol (9:1) GUVs demonstrated a gradual inward membrane remodelling, similar to pure POPC vesicles. Internal membrane tubules, intraluminal buds and daughter-vesicle-like compartments were seen during the course of starch hydrolysis. The addition of cholesterol did not significantly alter the general sequence of morphological alterations as shown in Figure 9. Similar inward budding and production paths of daughter vesicles are seen.

**Fig. 9.**
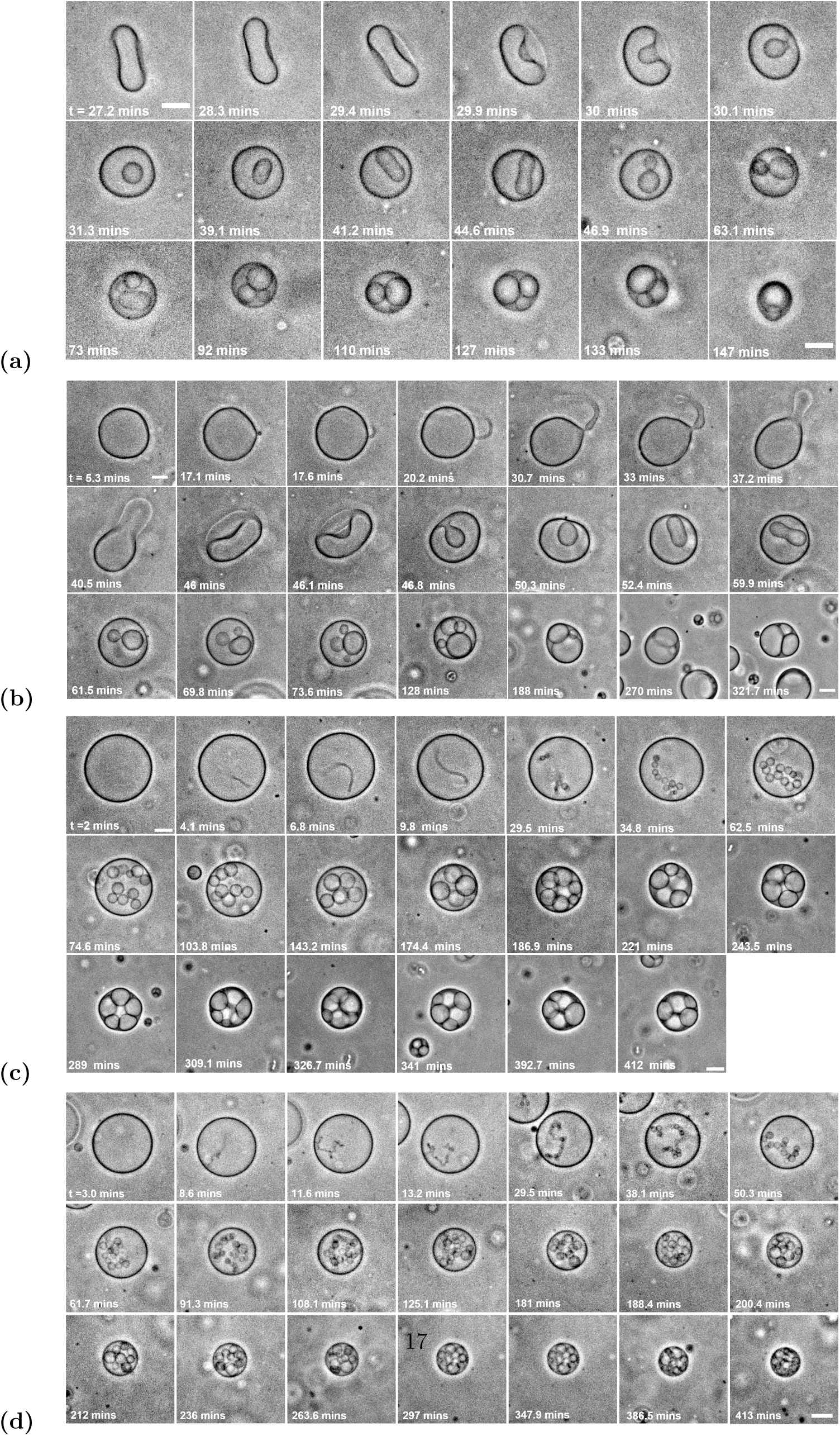
Time-lapse microscopy images showing morphology changes of POPC: Cholesterol (9:1) GUVs prepared in 0.5 mg/mL starch in the presence of increasing concentrations of *α*-amylase: (a) 1 *µ*M, (b) 1.5 *µ*M, (c) 2 *µ*M and (d) 2.5 *µ*M, Scale bar: 10 *µ*m for all images.

When the concentration of *α*-amylase exceeds 2.5 *µM*, a marked disruption in the structural integrity of the GUVs is observed (see Supporting Information, Figure S4). This disruption highlights a critical threshold above which the stability of the GUVs is compromised, emphasising the need for careful control of *alpha*-amylase concentration to maintain vesicle integrity in experimental and practical applications.

The extent of disruption was also found to depend on the initial size of the vesicles. Larger vesicles, particularly those with diameters greater than 25 *µm*, were able to survive even at 3 *µ*M *α*-amylase concentration, whereas the majority of smaller vesicles underwent complete disruption. These observations indicate that both enzyme concentration and vesicle size play important roles in determining the mechanical stability and survival of GUVs during enzymatic starch hydrolysis.

### 3.6 Alpha-Amylase Induced Remodeling of Phase-Separated GUVs

Salivary membranes are composed of a heterogeneous mixture of lipids rather than a single phospholipid species. To further increase the compositional complexity of the model membrane, we investigated ternary GUVs composed of DOPC:DPPC:Cholesterol (0.35:0.35:0.30). GUVs were prepared in the same 0.5 mg/mL starch solution and examined before and after the addition of externally supplied *α*-amylase. Under the experimental conditions used here, these GUVs exhibited lateral phase separation, providing a membrane system with distinct lipid domains. In the absence of *α*-amylase, the ternary GUVs displayed distinct membrane domains while maintaining an overall vesicular morphology. Following the addition of *α*-amylase, the GUVs exhibited both the pre-existing membrane domains and pronounced morphological remodeling, including inward budding and the formation of internal compartments as shown in Figure 10. Thus, enzymatic remodeling was observed in a laterally heterogeneous membrane in which membrane domains were already present.

**Fig. 10.**
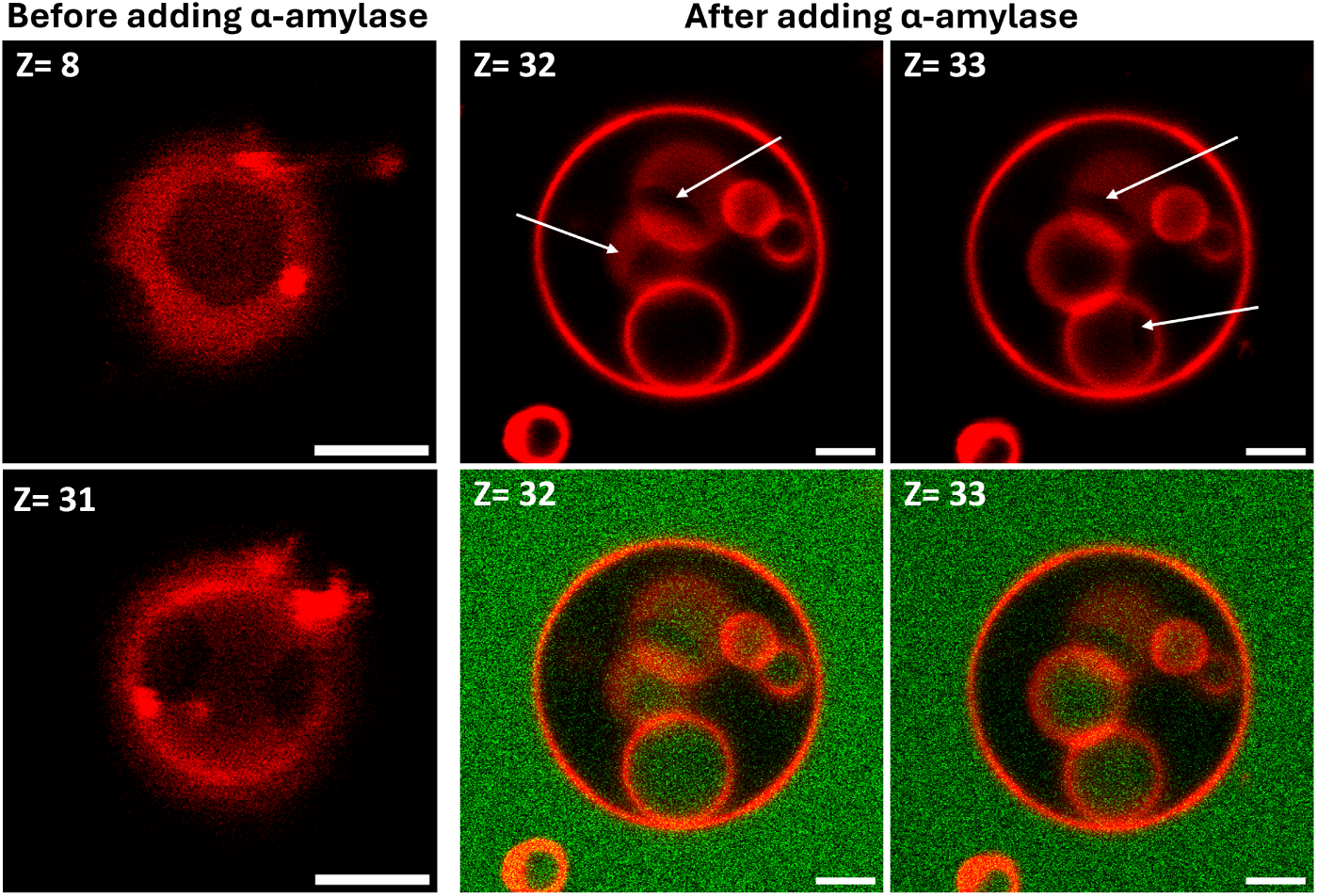
Confocal z-stack images of phase-separated ternary DOPC:DPPC:Cholesterol GUVs prepared in 0.5 mg/mL starch solution, before and after the addition of fluorescently labelled *α*-amylase at a concentration of 1 *µ*M on the outside.Before enzyme addition, the same GUV is shown at different z-positions, exhibiting phase-separated membrane domains without visible budding structures (left). Following the addition of *α*-amylase and 3 h incubation, a different GUV is shown at consecutive z-positions, exhibiting phase-separated membrane domains together with morphologically distinct budding/internal membrane structures (middle and right). Nile Red was added immediately before imaging. Arrows indicate the phase-separated domains and budding/internal membrane structures. The corresponding merged images show the spatial distribution of *α*-amylase fluorescence relative to the membrane morphology. The z-stack analysis was used to distinguish membrane-associated domains from three-dimensional budding/internal structures. Scale bar is 5 *µ*m applied to all the images.

### 3.7 Localization of *α*-Amylase during GUV Remodeling

The time-dependent morphological alterations in GUVs during starch hydrolysis were continuously observed using phase-contrast microscopy (see Figure 7). To get more insight into the interaction of *α*-amylase with the membrane and to find out whether the enzyme enters the vesicles, FITC-labelled *α*-amylase was monitored by confocal fluorescence microscopy. For these investigations, 2.5 *µ*M FITC-tagged *α*-amylase (green fluorescence) was added outside of the GUVs. The two types of vesicles studied were vesicles without internal budding and vesicles with inward bud development. In vesicles without budding, no green fluorescence was seen inside the vesicle lumen, showing that *α*-amylase does not permeate straight through the intact lipid bilayer, as shown in Figure 11.

**Fig. 11.**
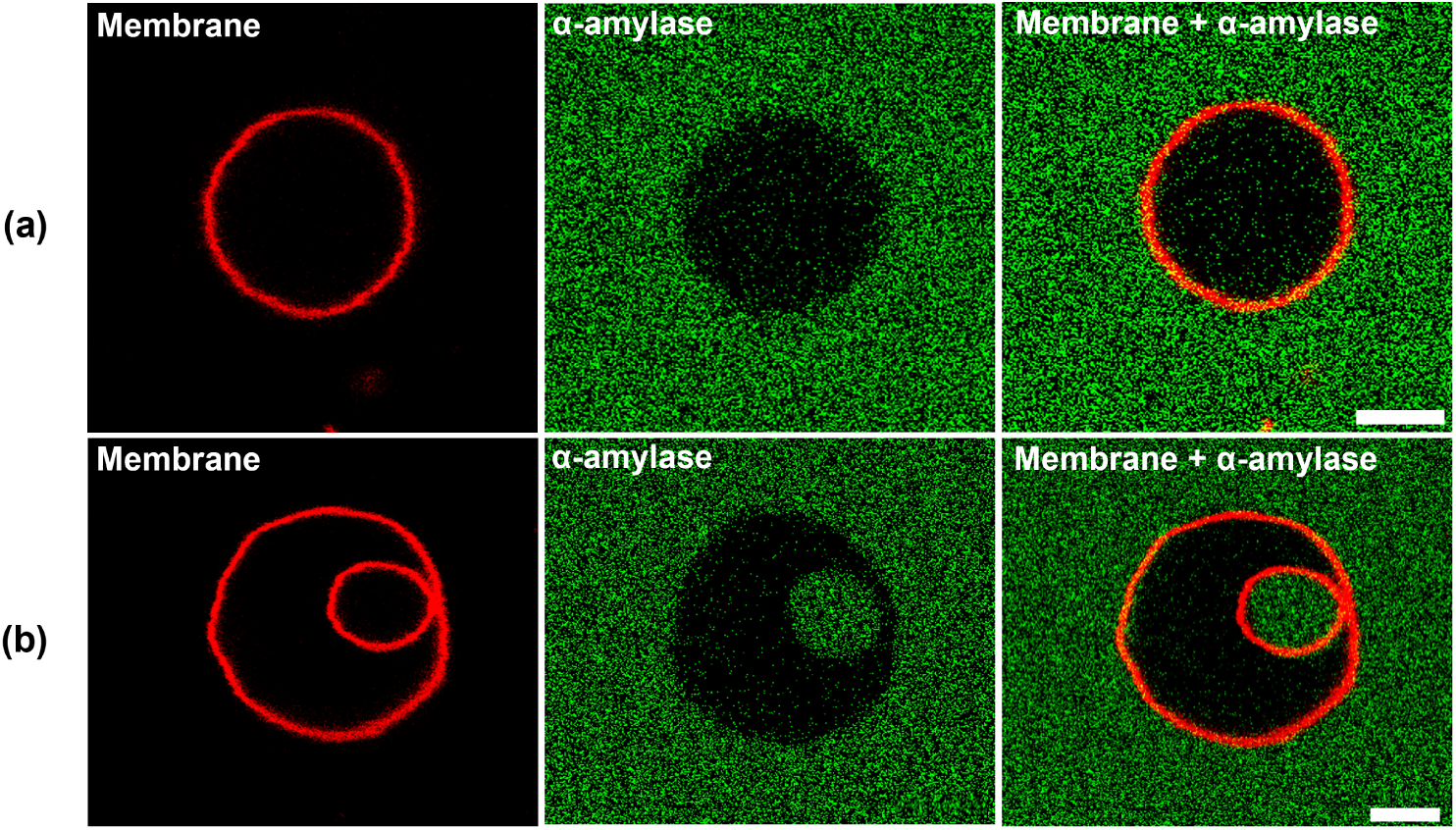
Confocal fluorescence images of POPC GUVs prepared in 0.5 mg/mL starch solution, with FITC-labelled *α*-amylase at a concentration of 2.5 *µ*M on the outside.The lipid membrane is labelled in red, while FITC-tagged *α*-amylase appears in green. (a) vesicle without membrane budding, with no green fluorescence in the vesicle lumen, suggesting that enzyme internalisation through the intact membrane is absent. (b) a vesicle with inward budding, where green fluorescence is found in the budded compartment, indicating that *α*-amylase is taken up by the budding-mediated membrane transport. Scale bar is 5 *µ*m applied to all the images.

### 3.8 Membrane Mechanical Response

The GUVs are fluid lipid membrane systems that are very soft and flexible. The deformation of a membrane, caused by collisions and the exchange of thermal or active energy with its surroundings, leads to the phenomenon known as membrane flickering or fluctuations [35]. Analysis of these thermally driven fluctuations provides a reliable method for determining the membrane bending rigidity (*κ*), which represents the energetic cost required to deform or bend the membrane.

In this study, fluctuation spectroscopy [32, 36] was performed for POPC GUVs suspended in starch solution in the absence of *α*-amylase. The bending rigidity (*κ*) was calculated from the contour fluctuations of ten individual vesicles, and the corresponding plot is shown in Figure 12. The obtained bending rigidity value is summarised in Table 3, where it is compared with previously reported values for POPC bilayers measured under different surrounding media and experimental conditions.

**Fig. 12.**
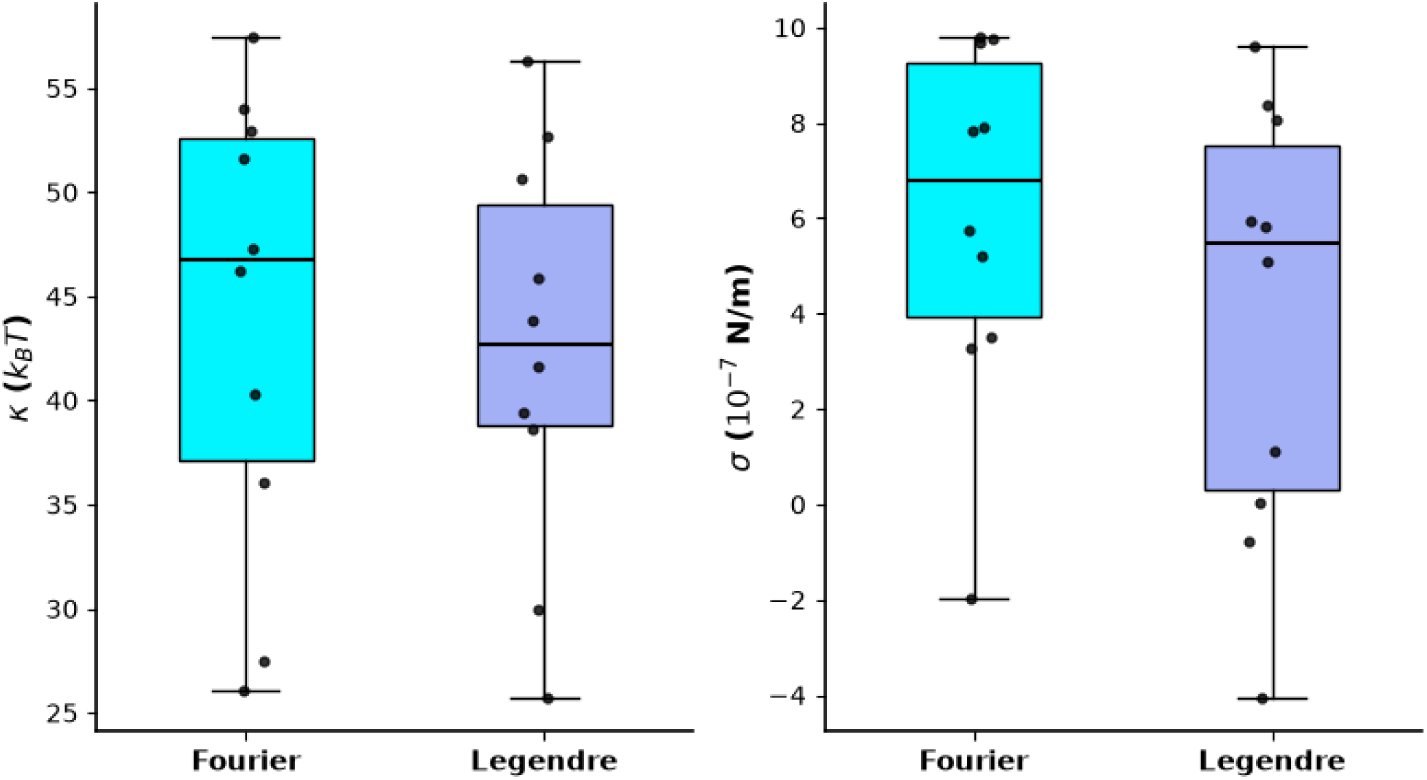
Bending rigidity and membrane tension data for the POPC GUVs prepared in 0.5 mg/mL starch solution in the absence of *α*-amylase. The Fourier and the Legendre modes analysis was performed for vesicles, and the resulting *κ* and membrane tension values are shown.

**Table 3.**
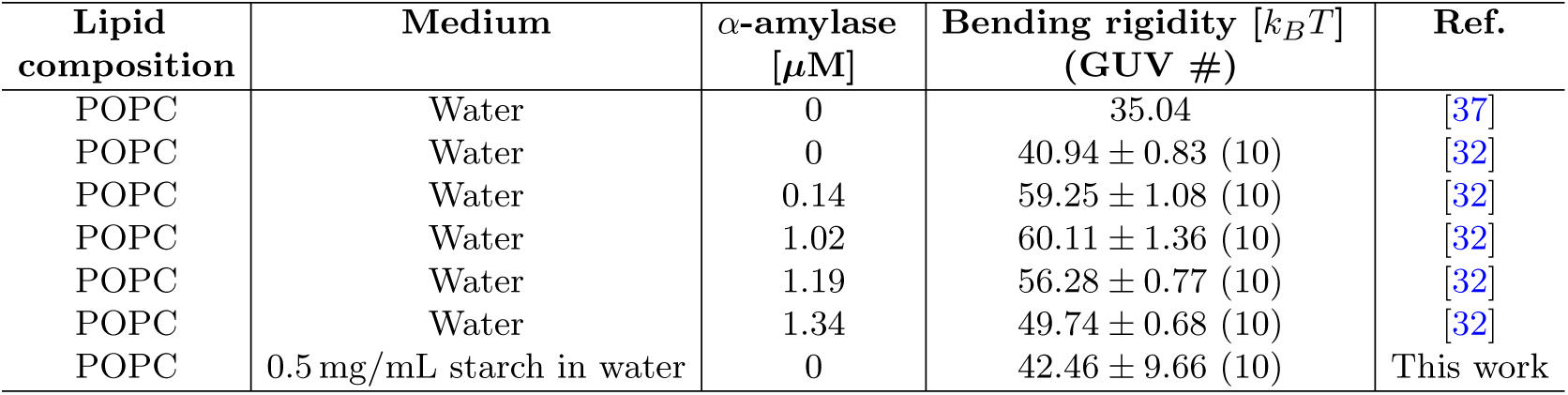
Summary of bending rigidity (*κ*) values of POPC membranes obtained from fluctuation spectroscopy under different conditions.

| **Lipid composition** | **Medium** | **α-amylase [µM]** | **Bending rigidity [*k*<sub>*B*</sub>*T*] (GUV #)** | **Ref.** |
|---|---|---|---|---|
| POPC | Water | 0 | 35.04 | [37] |
| POPC | Water | 0 | 40.94 ± 0.83 (10) | [32] |
| POPC | Water | 0.14 | 59.25 ± 1.08 (10) | [32] |
| POPC | Water | 1.02 | 60.11 ± 1.36 (10) | [32] |
| POPC | Water | 1.19 | 56.28 ± 0.77 (10) | [32] |
| POPC | Water | 1.34 | 49.74 ± 0.68 (10) | [32] |
| POPC | 0.5 mg/mL starch in water | 0 | 42.46 ± 9.66 (10) | This work |

The value measured for POPC GUVs suspended in a starch solution of concentration 0.5 mg/mL in the absence of *α*-amylase was

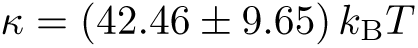

This value is slightly higher than the value we previously found for POPC vesicles in pure water *κ* = 40.94 *±* 0.83 *k*_B_*T* (see Table 3). The increase in bending rigidity indicates that the presence of starch in the surrounding medium makes the membrane more stiffer to bending deformations.

This behaviour is likely due to the influence of starch molecules on the physical environment surrounding the membrane. Through excluded-volume effects, increased solution viscosity, or weak interactions with the membrane surface, starch can suppress membrane undulations and effectively increase the energy required for membrane bending. As a result, the POPC bilayer exhibits enhanced mechanical stability in starch-containing solutions.

A comparison of the starch-only condition with samples containing *α*-amylase shows that starch alone induces a substantial stiffening effect. The addition of *α*-amylase further modifies the membrane rigidity, with *κ* reaching a maximum value of approximately 60 *k*_B_*T* at intermediate enzyme concentrations of 0.14–1.02 *µ*M, before decreasing at higher concentrations. This trend suggests that enzymatic modification of starch influences membrane mechanics. This baseline characterisation establishes the mechanical state of the membrane prior to enzymatic activity and provides an important reference for understanding the membrane remodelling observed upon *α*-amylase-mediated starch hydrolysis. The analysis using Fourier and Legendre modes are described in the our previous work [32]. The individual values of the bending rigidity is shown in the Fig. 13. We have from the FOURIER analysis, *κ* = (43.95 *±* 11.08) *k_B_T*, *σ* = (6.08 *±* 3.75) *N/m* and for the LEGENDRE analysis, *κ* = (42.46 *±* 9.66) *k_B_T* and *σ* = 3.92 *±* 4.56 *N/m*.

**Fig. 13.**
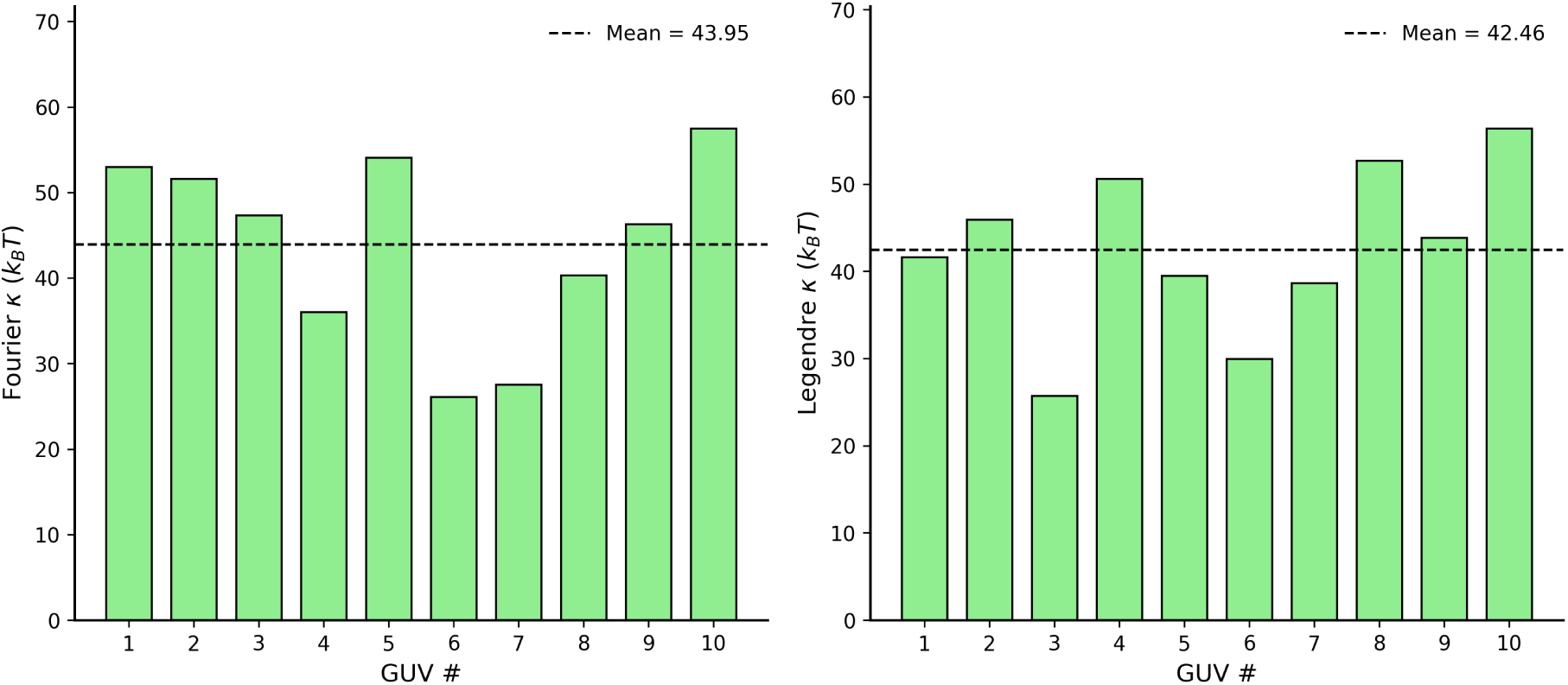
Bending rigidity and membrane tension data for the POPC GUVs prepared in 0.5 mg/mL starch solution in the absence of *α*-amylase. The Fourier and the Legendre modes analysis was performed for vesicles, and the resulting *κ* and membrane tension values are shown.

## 4 Conclusion

In this study, we demonstrate that external enzymatic activity can serve as an effective physicochemical trigger for membrane remodelling in giant unilamellar vesicles (GUVs). By preparing vesicles under initially symmetric osmotic conditions in starch solution and subsequently introducing *α*-amylase into the external medium, we establish a minimal model system in which biochemical reactions directly translate into mechanical membrane responses.

Analysis of the potato starch utilised in this work by UV-Visible iodine-binding spectroscopy showed that the starch was composed of 26% amylose and 75% amylopectin, confirming the dominance of amylopectin. The dominance of amylopectin is particularly important because the highly branched structure of amylopectin offers many sites for accessible cleavage by *α*-amylase, allowing effective enzymatic breakdown.

Quantitative DNS analysis revealed time-and concentration-dependent accumulation of reducing sugars during starch hydrolysis. The progressive increase in low-molecular-weight hydrolysis products generated a measurable osmotic pressure difference across the lipid bilayer, reaching approximately 4.3 kPa at the highest enzyme concentration investigated. Since the DNS assay measurements were performed using a 15-fold diluted reaction mixture (200 *µ*L sample diluted to 3 mL), correspond to the diluted system. In the actual GUV experiments, where the starch concentration was 0.5 mg/mL, the corresponding osmotic pressure is expected to be approximately 15 times higher than the values presented here.

This externally generated osmotic imbalance drives water efflux from the vesicle interior, reducing vesicle volume while conserving membrane area. The resulting decrease in reduced volume promotes curvature instability, leading to inward budding and the formation of internal compartments, as observed by time-lapse phase-contrast microscopy.

The type of hydrolysis products was determined by HPLC-RID analysis using glucose and malto-oligosaccharide standards. The chromatograms demonstrated the synthesis of glucose, maltose, maltotriose, maltotetraose, maltopentaose, maltohexaose and maltoheptaose with maltose and maltotriose as the major products. Product distributions were similar for all enzyme concentrations examined. Peak intensities increased with increasing enzyme concentration, indicating more soluble carbohydrates produced with higher hydrolytic activity.

These combined results indicate that *α*-amylase is able to degrade potato starch into a mixture of osmotically active sugars and malto-oligosaccharides. The accumulation of these soluble hydrolysis products slowly increases the osmolarity of the extracellular media surrounding the vesicles, leading to the establishment of osmotic asymmetry across the membrane. This biochemical framework provides the quantitative basis for the understanding of the processes of membrane deformation, generation of curvature, vesicle deflation and budding.

In addition to the specific starch–*α*-amylase system studied here, our work uncovers a general mechanism for external biochemical reactions to control membrane shape transformations via osmotic coupling. Such enzyme-driven remodelling offers a simple yet powerful strategy for inducing dynamic structural changes in artificial cells. It may provide insight into primitive protocellular processes in which metabolic activity and membrane mechanics were inherently coupled.

## Acknowledgements

This research was conducted within the Indian Institute of Science Education and Research Mohali and supported by the Ramalingaswami grant of DBT (DBT/RLF/Re-entry/06/2020), DST/INT/DAAD/P-1 6/2023 (G) and IC-12025(22)/1/2023-ICD-DBT. PhD fellowship for R.K. is supported by IISER Mohali.

## Declarations

The authors declare no competing interests. R.K. conducted experiments and prepared experimental figures, wrote the manuscript. S.N. handled HPLC experiments and prepared plots. A.S. contributed to the bending rigidity analysis. D.S. participated in designing the HPLC experiments, and analyzing the HPLC data. T.B. participated in designing the experiments, collecting and analyzing the data (except HPLC), interpreting the results, and preparing the manuscript. All authors reviewed and approved the final manuscript.

## Notes

### Competing Interest Statement

The authors have declared no competing interest.

